# Dose-dependent enhancement of motion direction discrimination with transcranial magnetic stimulation of visual cortex

**DOI:** 10.1101/2020.06.14.151118

**Authors:** Olga Lucia Gamboa Arana, Hannah Palmer, Moritz Dannhauer, Connor Hile, Sicong Liu, Rena Hamdan, Alexandra Brito, Roberto Cabeza, Simon W. Davis, Angel V. Peterchev, Marc A. Sommer, Lawrence G. Appelbaum

## Abstract

Despite the widespread use of transcranial magnetic stimulation (TMS) in research and clinical care, the underlying mechanisms-of-actions that mediate modulatory effects remain poorly understood. To fill this gap, we studied dose–response functions of TMS for modulation of visual processing. Our approach combined electroencephalography (EEG) with application of single pulse TMS to visual cortex as participants performed a motion perception task. During participants’ first visit, motion coherence thresholds, 64-channel visual evoked potentials (VEPs), and TMS resting motor thresholds (RMT) were measured. In second and third visits, single pulse TMS was delivered 30 ms before the onset of motion or at the onset latency of the N2 VEP component derived from the first session. TMS was delivered at 0%, 80%, 100%, or 120% of RMT over the site of N2 peak activity, or at 120% over vertex. Behavioral results demonstrated a significant main effect of TMS timing on accuracy, with better performance when TMS was applied at N2-Onset timing versus Pre-Onset, as well as a significant interaction, indicating that 80% intensity produced higher accuracy than other conditions. TMS effects on VEPs showed reduced amplitudes in the 80% Pre-Onset condition, an increase for the 120% N2-Onset condition, and monotonic amplitude scaling with stimulation intensity. The N2 component was not affected by TMS. These findings reveal dose–response relationships between intensity and timing of TMS on visual perception and electrophysiological brain activity, generally indicating greater facilitation at stimulation intensities below RMT.

## 1. Introduction

Transcranial magnetic stimulation (TMS) has become a valuable treatment option for a host of psychiatric and neurological disorders and a useful tool in the study of the psychophysiology of human cognition. The underlying mechanisms-of-action that lead to these effects, however, are relatively poorly understood, hence the strong emphasis on filling this gap through the ongoing National Institute of Health, BRAIN Initiative research priorities (NIH, 2019). While a rich literature of studies now offer characterization of transient induced, and plastic long-term, effects of TMS in the motor system (Hallett et al., 2017; Pascual-Leone et al., 1995), systematic dose–response characterization in the visual system is lacking.

TMS to the visual system can yield non-retinal perceptions, referred to as “phosphenes” for static flashes or “mophenes” if perceptual motion is induced (Pascual-Leone & Walsh, 2001; Schaeffner & Welchman, 2017), as well as modulation of perception from retinal input (de Graaf, Koivisto, Jacobs, & Sack, 2014). Notably, single-pulse TMS (spTMS) and paired-pulse TMS (ppTMS) of the visual cortex have been widely reported to induce changes in the perception of visual motion (Bosco, Carrozzo, & Lacquaniti, 2008; Grasso et al., 2018; Laycock, Crewther, Fitzgerald, & Crewther, 2007; Silvanto, Lavie, & Walsh, 2005; Vetter, Grosbras, & Muckli, 2015). Across a number of studies, TMS to motion sensitive cortex has been shown to influence speed (Matthews, Luber, Qian, & Lisanby, 2001) and direction sensitivity (Campana, Cowey, & Walsh, 2002), as well as perception of biological motion (Mather, Battaglini, & Campana, 2016). Despite this accumulation of literature, the relationship between the reported physiological response and the degree of behavioral engagement across this literature is highly variable, and the dose-response relationships between TMS and neurophysiology have yet to be established.

Studies of motion perception are particularly well-suited for exploring mechanisms of TMS due to the superficial location of motion sensitive cortex and its well-characterized spatiotemporal progression of electroencephalographic (EEG) activation described by the P1/N2/P3 Visual Evoked Potential (VEP) complex. In this progression, it is regarded that the initial P1 reflects pattern-related activity of the parvo-cellular subsystem, while the subsequent N2 has been associated with motion perceptual sensitivity (Bach & Ullrich, 1997; Kuba, Kubová, Kremláček, & Langrová, 2007; Martin, Huxlin, & Kavcic, 2010) and is localized to the direction-selective area V5 of the extra-striate visual cortex (Pazo-Álvarez, Amenedo, Lórenzo-López, & Cadaveira, 2004). In lateralized visual attention tasks, this response has particularly been characterized as pre-attentive in the early <200 ms phase, and specific to spatial attention after about 200 ms, as observed in the widely-reported N2pc component (Clark, Appelbaum, van den Berg, Mitroff, & Woldorff, 2015; Luck & Hillyard, 1990). Lastly, in tasks where response selection is made on motion stimuli, a central positive P3 component is frequently observed around 300 ms (Kuba, Kremláček, & Kubová, 1998; Kubová, Kremláček, Szanyi, Chlubnová, & Kuba, 2002) that is thought to reflect attentional allocation to the stimuli (Duncan-Johnson & Donchin, 1982). While other thalamo-cortical pathways also contribute to visual motion perception, these VEPs offer specific testable neural markers that can be characterized to infer dose–response properties of TMS

The present study builds on these two bodies of literature, TMS modulation of motion perception and motion-induced VEPs, to characterize dose–response functions of spTMS in the visual cortex. Our approach was to measure concurrent TMS-EEG during an individually calibrated, dot-motion, direction-discrimination task, previously developed by our group to test ppTMS effects (Gamboa, 2019). For this purpose, we used a three-visit study design consisting of an initial dose-finding session to derive individualized motion coherence thresholds and stimulation parameters based on the onset of the N2 VEP component (“N2-Onset”), followed by two dose-testing sessions during which spTMS was delivered according to the spatial, temporal, and intensity parameters derived from the first session. During the dose-testing sessions, spTMS was delivered at one of two different timings, either 30 ms before the onset of motion (“Pre-Onset”, based on the observation that TMS to V5 around this latency can disrupt motion perception (Laycock et al., 2007; Sack, Kohler, Linden, Goebel, & Muckli, 2006)), or at the individualized N2-Onset latency from Session 1. During each of these two sessions, participants performed eight blocks of trials with intermixed pulse intensities at 0%, 80%, 100%, and 120% of RMT delivered over the hotspot of the N2-Onset component, as well as two separate blocks of trials at 120% RMT at a vertex, control location. As such, this study tested the effects of spTMS over intensities that have previously been reported to induce facilitatory and inhibitory perceptual effects (Luber et al., 2020; Silvanto, Bona, & Cattaneo, 2017), but at different timings relative to the motion onset and at different cortical targets. The overall goal was to map the behavioral and evoked EEG dose–response functions within the constructs of individualized spatiotemporal targeting with the expectation that spTMS delivered at the onset of the N2 component would disrupt motion processes in the brain and lead to monotonic effects across different stimulation intensities.

## 2. Methods

### 2.1. Participants

Twenty-four healthy volunteers (15 females, mean age = 23, SD = 2.55) enrolled in this 3-visit study. All participants were self-reported right-handed, had normal or corrected-to-normal vision, and were screened for contraindications to TMS (Rossi et al., 2009). Exclusion criteria included a history of neurological or psychiatric disease and/ or use of psychoactive medication, a personal history of head trauma with loss of consciousness or family history of epilepsy or seizures. Informed consent was obtained for each participant after explanation of study requirements under an experimental protocol approved by the Duke University Institutional Review Board (Pro00082433). Participants were compensated $20 per hour for their time.

### 2.2. Experimental Design and Equipment

This study consisted of three sessions, each lasting 2 to 3 hours, performed on separate visits within a three-week span. Twenty-one participants completed the first dose-finding session, with 18 and 17 completers for the Pre-Onset and N2-Onset sessions, respectively. In total, 15 of the participants completed all three sessions, while the remaining participants only completed one or two of the sessions due to scheduling conflicts, illness, or equipment difficulty.

All stimuli were generated using Matlab (Mathworks, Natick, MA, USA) and the Psychtoolbox extension (Brainard, 1997) and presented on a 21-inch monitor with 60 Hz refresh and 1920 × 1050 screen resolution. EEG was acquired on all three visits using a 64-channel actiCHamp EEG system (BrainProducts, Munich, Deutschland) and TMS compatible actiCAP with slim active electrodes arranged according to an equidistant montage. The EEG signal was digitized at a sampling rate of 5 kHz and the online reference electrode was located at FCz. TMS was delivered using a MagPro R30 stimulator connected to a Cool-B65 figure-of-eight coil (MagVenture, Farum, Denmark). Synchronization between the stimulation computer cand TMS machine was done through an Arduino board that allowed a temporal precision between systems that was confirmed to be approximately 1 ms.

A Brainsight (Rogue Research, Canada) frameless stereotactic neuronavigation system was used to monitor TMS coil position on the participant’s head so that the stimulation location was kept as constant as possible. For this, coil trackers were attached to the coil and the participant’s head, which was registered to a standard head model using anatomical landmarks. A coil holder and a chin rest were also used to maintain stable coil and head positioning. In addition, a spacer was used to minimize contact with the electrodes and associated artifact (Ruddy et al., 2018), while a thin foam sheet (∼1 mm) was added to minimize bone conduction of the TMS sound and scalp sensations caused by mechanical vibration of the coil. Participants wore earplugs for hearing protection and mitigation of auditory activation by the TMS pulse clicks. Two experimenters were always present in the room, to maintain coil positioning and supervise EEG data quality.

#### 2.2.1. Session 1: Dose-Finding

During the first session, eligibility was assessed and consent was obtained. Following this, the 64-channel EEG cap was fitted to the participant’s head and after applying gel the Resting Motor Threshold (RMT) was acquired, taking about 15 minutes according to the procedures described below. This allowed for estimations of stimulation intensity calculation according to RMT with equivalent distancing between the head and coil as used in the dose-testing sessions. Participants then performed 20 minutes of practice with the motion task (**Figure 1A**) while EEG impedances were being adjusted and recordings were prepared. After obtaining clean EEG signals, with impedances below 5 kΩ, the participant completed four, 8-minute runs of the motion discrimination task.

**Figure 1:**
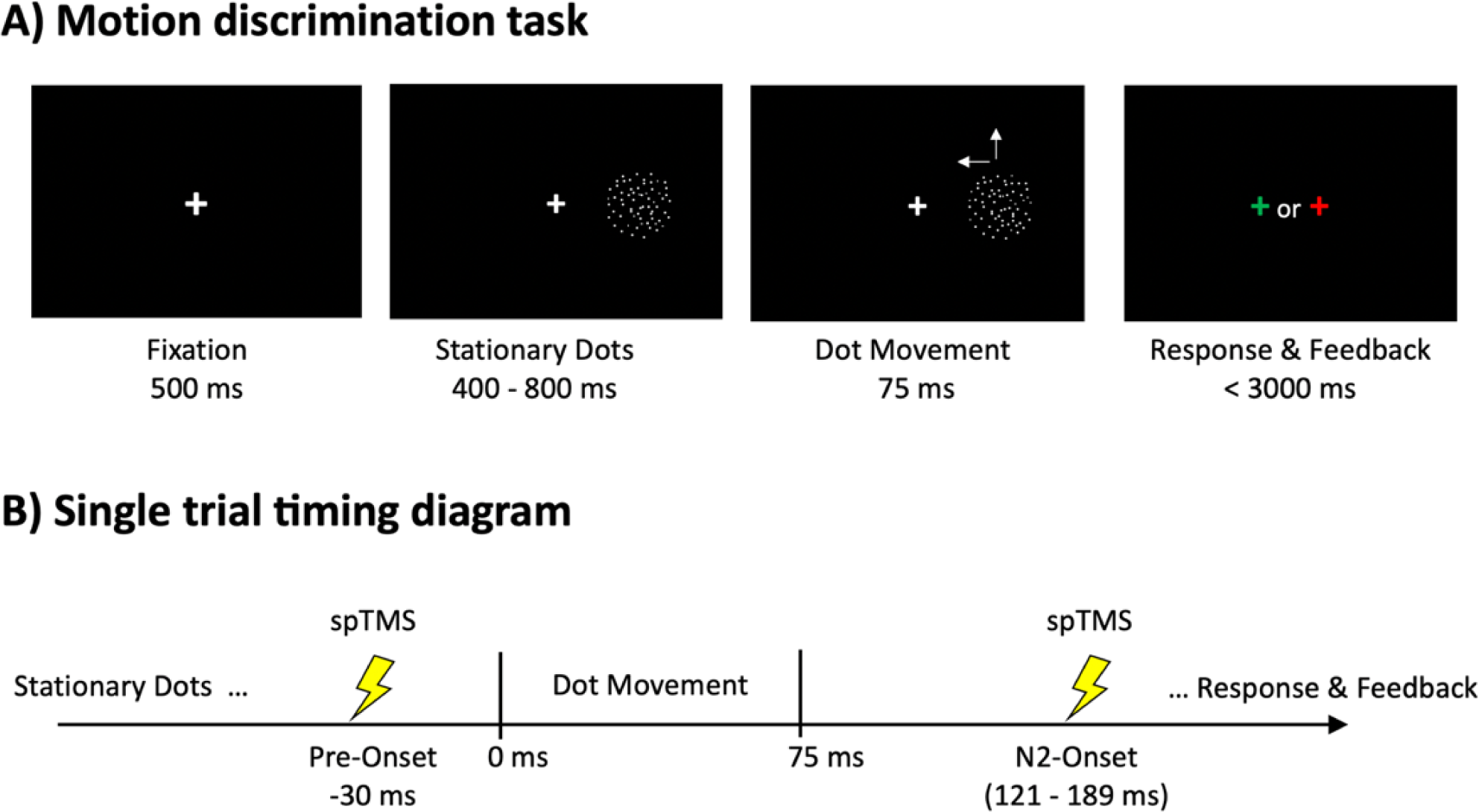
Schematic illustrations of A) motion direction discrimination task showing four example frames and the duration of each sequence, and B) schematic of TMS (lightning bolts) relative to dot motion.

##### Resting Motor Threshold (RMT)

RMT was defined as the lowest intensity required to elicit a motor evoked potential (MEP) of 50 μV peak-to-peak when the muscle was at rest (Conforto, Z’Graggen, Kohl, Rösler, & Kaelin-Lang, 2004). Surface electromyogram was recorded from the right first dorsal interosseous muscle by using disposable electrodes (Covidien/Kendall, 133 Foam ECG Electrodes, Mansfield, USA) in a belly tendon montage. After establishing a reliable stimulation site according to the visual-twitch method, RMT was determined by a probabilistic threshold-hunting method using the Motor Threshold Assessment Tool software (http://www.clinicalresearcher.org/software.html).

##### Motion Direction Discrimination Task and Visual Evoked Potential

A common dot motion discrimination task was employed across all sessions of this experiment. In the first ‘dose-finding’ session of the study, the coherence of dot motion varied randomly from trial to trial, allowing for the estimation of each individual’s threshold coherence value, which was then fixed to this level for each participant in sessions 2 and 3.

In this task, each trial initiated with a white fixation cross appearing for 500 ms, after which two fields of static white dots appear within 12.6° circular windows centered 7.6° to the left and right of fixation. These dots remained static for a variable duration between 400 and 800 ms, after which the dots moved briefly for 75 ms at a speed of 15.5 °/s. The dots in left field moved incoherently in random directions while the dots in the right field contained some level of directional coherence, determined by the staircase procedures. On each trial a dot coherence was randomly selected to be 0, 20, 40, 60, 80, or 100%. Following the motion, participants had 2 seconds to indicate with a button press if the direction of motion was to the left or upwards. After each answer, feedback was provided with a green fixation cross for correct responses and red for incorrect responses that was presented for 1 s. The viewing distance (eye-to-screen center) for each participant was kept constant at approximately 56 cm and all the participants were instructed to perform the task as accurately as possible within the 2 second allotted time.

At the end of the session, a generalized linear model was applied to fit a sigmoid function to the assigned (+1/−1 due to dot motion direction) trial coherences and correct/incorrect responses to determine the 75% accuracy point on the coherence psychometric curve. If 75% accuracy could not be achieved, then 100% dot motion coherence was used. As described in greater detail in section 2.3.2, concurrent EEG was collected and analyzed for VEP results. As described in Section 3.1, these analyses led to characterization of an N2 onset latency and location that was used for dosing in Sessions 2 and 3.

#### 2.2.2. Sessions 2 and 3: Dose-Testing

During Sessions 2 and 3, participants performed the same motion task with dot stimuli presented only at the threshold coherence level determined from Session 1. These sessions differed only in the latency at which TMS was applied relative to the onset of the motion on each trial. As illustrated in **Figure 1B**, biphasic spTMS was applied either 30 ms before the onset of motion (Pre-Onset) or at the onset latency of the N2 component (N2-Onset) derived from Session 1. The V5 target was defined as the topographic location showing the most robust N2 response identified from the current source density maps (CSD) corresponding to the N2 sink topography (see right column of **Table 1**). Consistent with what is known about motion sensitive cortex (Silvanto et al., 2005), this topographic localization pointed to electrodes over the left occipital cortex, as the regions displaying maximal activity (referred to here as V5). The center of the coil was placed tangentially to this stimulation site with the handle pointing towards the right hemisphere.

**Table 1:** Experimental parameters collected in Session 1. Stimulation intensity at RMT, motion coherence thresholds, stimulation times and closest stimulated channels (See **Figure 1A** for locations) for the closest EEG electrodes relative to the position of stimulation in Sessions 2 and 3. The coil was positioned approximately over the center of mass surrounding these electrodes, based on the N2 topography in visit 1, as guided by BrainSight neuronavigation.

|  | **RMT**<br>(% MSO) | **Percent Dot**<br>**Coherence** | **Stimulation**<br>**times (ms)** | **Closest stimulated**<br>**channels** |
| --- | --- | --- | --- | --- |
| **S1** | 54 | 37 | 184 | 18-20 |
| **S2** | 57 | 74 | 160 | 20-24-21 |
| **S3** | 81 | 100 | 161 | 20-24-25 |
| **S4** | 57 | 75 | 143 | 23-24 |
| **S5** | 83 | 92 | 170 | 20-24 |
| **S6** | 55 | 100 | 119 | 25-51 |
| **S7** | 83 | 100 | 156 | 21-24 |
| **S8** | 71 | 100 | 166 | 24-25 |
| **S9** | 64 | 100 | 129 | 20-21-24-25 |
| **S10** | 64 | 100 | 189 | 20-24-25 |
| **S11** | 62 | 82 | 151 | 24-25 |
| **S12** | 65 | 100 | 158 | 20-25 |
| **S13** | 82 | 100 | 121 | 20 |
| **S14** | 55 | 100 | 152 | 20-24 |
| **S15** | 80 | 100 | 153 | 24-20-21 |
| **S16** | 58 | 50 | 148 | 25-20-24 |
| **S17** | 52 | 100 | 141 | 25-20-24 |
| **S18** | 50 | 76 | 163 | 20-21-24 |
| **S19** | 63 | 100 | 125 | 25-20 |
| **S20** | 77 | 100 | 174 | 24-23-21 |
| **S21** | 44 | 84 | 148 | 24-25-20 |

On both of these sessions the procedures were identical and began with the participant practicing the motion task as the cap was placed, gel applied, and impedances checked and adjusted. After a clean EEG signal was achieved with impedances below 5 kΩ, the participant began eight 4-minute blocks of 56 trials each while receiving spTMS. In six of the eight blocks, TMS was delivered to the V5 target channel at intensities of 0%, 80%, 100% or 120% RMT, controlled remotely using a customized function from MAGIC Matlab toolbox (Saatlou et al., 2018). These intensities were consistently delivered for all trials within a block and changed according to a random sequence created for each block at the beginning of the session. The two remaining blocks were assigned to stimulation of the vertex during task performance at 120% of RMT (112 trials). Stimulation over vertex was performed as a control condition. Vertex was defined as the scalp location corresponding to Cz in the 10–20 system. The coil handle was parallel to the midline pointing backwards. Two blocks of stimulation delivered over vertex were randomly distributed throughout the eight total blocks, as determined by a ordering calculation done at the start of each session.

### 2.3. Analyses

#### 2.3.1. Behavioral data analyses

One-way ANOVA were performed to evaluate the effect of dot coherence in the initial dose-finding session. To evaluate experimental effects in the dose-testing sessions changes in accuracy were examined by a repeated measure ANOVA (rmANOVA) with factors Stimulation Condition (0%, 80%, 100%, and 120% RMT at V5 and 120% RMT at vertex) and TMS Timing (Pre-Onset and N2-Onset). Additionally, to provide a more sensitive assessment of the interaction between TMS at different intensities and brain dynamics, accuracy data were also analyzed as the difference of each Stimulation Condition, minus the 0% RMT condition. This condition presents the same visual stimuli but no TMS pulse or associated sounds and sensation, making it an effective baseline to normalize ongoing tonic activity. A 2 × 4 rmANOVA was performed over these differences. Post hoc pairwise comparisons were tested using Tukey’s HSD method. A Shapiro-Wilks test was used to verify the assumption of normality while Mauchly’s test evaluated the assumption of sphericity. A Greenhouse-Geisser correction was adopted when the sphericity assumption was violated. Statistical tests were performed using SPSS 25.0 (SPSS Inc, Chicago) and *p* values ≤ .05 were considered statistically significant.

#### 2.3.2. EEG data processing and analysis

EEG data were preprocessed and analyzed offline using Brain Vision Analyzer (Brain Products, Inc.) and Matlab (Mathworks, Natick, MA, USA), in a manner modeled after Rogasch et al. (2014). Data were down sampled to 1000 Hz, TMS pulses were removed via linear interpolation spanning from 10 ms before to 25 ms after the pulse. Before applying Independent Component Analysis (ICA) to identify and discard artifactual components such as eye blinks and muscle activity, the EEG signal was bandpass filtered using a zero-phase shift Butterworth filter (0.1–45 Hz) and visually inspected to remove contamination. ICA-based artifact removal was then performed, leading to an average of 10.4 removed components per participant, with no significant difference in number of components removed across the TMS Timing sessions. Bad electrodes were interpolated before segmentation and baseline correction was performed from −250 to −50 ms for Pre-Onset and -200 to 0 ms for the N2-Onset averages. The final analyzed segments from −200 to 1000 ms were re-referenced to the common average across all channels.

VEPs for both the Pre-Onset and N2-Onset sessions were calculated by averaging artifact–removed trials in each channel. This led to an average of 78 trials per condition, per participant when stimulating at V5, and 103 trials per participant in the vertex stimulation condition. In order to investigate the electrophysiological correlates of visual motion perception, VEP analyses focused on mean activity in two multi-channel, regions-of-interest. As shown in **Figure 2D**, the V5 ROI comprised the channels 20, 21, 24 and 25, which surrounded the site of stimulation for all subjects. The Central Parietal (CP) ROI consisted of channels 1, 4, 18, 19, 33 and 34, centered over midline parietal cortex, in alignment with previously reported visual motion P3 effects (Kuba et al., 1998; Kubová et al., 2002). Because of the biphasic morphology of the N2 component in this study (See **Figure 2B**) and because stimulation in the N2-Onset timing disrupted the first phase of this component, the N2 was examined separately in an early (130–200 ms) and late (230–340 ms) latency window. Mean P3 amplitude was calculated between 250 and 550 ms.

**Figure 2.**
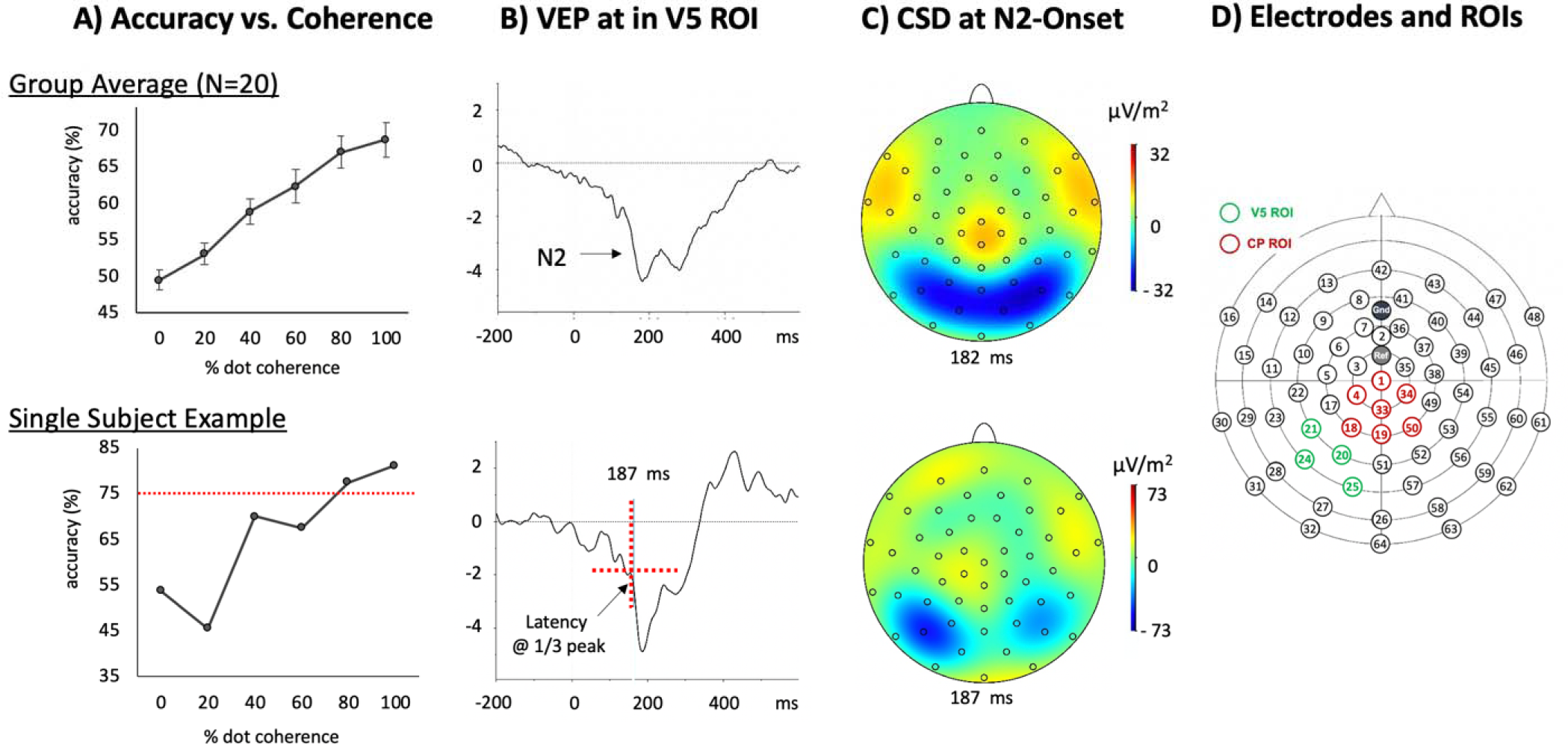
Behavior and VEP results from Session 1 shown on top for the 20-participant group average and on the bottom for a single subject. A) Coherence versus accuracy plots. B) VEP to motion onset for correctly reported trials at V5 ROI shows characteristic N2 component. C) Current source density topographic map at calculated N2-Onset. D) Electrode map of 64-channel EEG with V5 ROI (green) and Central Parietal ROI (CP; red). Vertex is channel 1.

Two participants were eliminated from the final ‘dose-testing’ analyses because of high levels of (predominantly blink) artifact in their N2-Onset sessions. As such, the final analyzed sample consisted of 18 participants with clean Pre-Onset data and 15 participants with clean N2-Onset data. Fourteen of these participants completed both TMS Timing sessions.

In order to focus analyses on the electrophysiological dose–response produced by different intensities, the following analyses compared the normalized, mean component amplitudes, subtracting the 0% (no stimulation) condition from each of the other conditions. These analyses were performed separately for the two TMS Timing sessions, allowing comparisons of components that would be obscured due to the different timing of the TMS pulse. For the Pre-Onset Timing, analyses addressed the early N2, late N2, and P3 components. In the N2-Onset Timing, analyses addressed the late N2 and P3 components. In addition, see **Supplement 1** for description of P1 component and reference sample.

## 3. Results

### 3.1. Session 1: Dose-Finding

#### 3.1.1. RMT Results

RMT was successfully derived for all participants through convergence of the adaptive Parameter Estimation by Sequential Testing (PEST) procedure (Borckardt, Nahas, Koola, & George, 2006). On average 63.6% of Maximum Stimulator Output (MSO) was required to induce the target EMG response at RMT with a range of 44% to 83%, as reported in **Table 1**.

#### 3.1.2. Accuracy and VEP Results

As illustrated in **Figure 2A**, participants performed the task at chance in trials that had no coherent movement (0%), with a monotonic increase in accuracy for higher coherence stimuli. One-way ANOVA confirmed a main effect of coherence (*F*[2.869,60.249] = 30.472, *p* < .001, *η*^2^ = 0.592) with an average dot coherence of 91.7% [range 34% – 100%] at threshold.

VEPs time locked to the onset of motion showed a waveform morphology (**Figure 2B** and **2C**) dominated by a bilateral negative posterior distribution peaking around 200ms, followed by a positive-going central-parietal deflection around 300 ms, suggestive of the widely reported N2 and P3 ERP components (Bach & Ullrich, 1997; Kuba et al., 2007; Martin et al., 2010). The N2 component was identifiable in all individual participant’s and was used to derive the N2-Onset stimulation timing and location. VEPs were processed to calculate the latency at 1/3 max amplitude of the N2 component at its peak location. These locations were always at left occipital channels (20, 21, 24, and 25), illustrated by the green labels in **Figure 2D**. Similar to previous studies using near-threshold motion onset (Kuba et al., 2007; Martin et al., 2010), the VEP here also did not contain a prominent P1 component (Bach & Ullrich, 1997; Kuba et al., 2007; Martin et al., 2010). The P1 response has been previously described as a pattern-sensitive parvocellular-driven component, which may have been absent here due to the 500 ms separation of the dot appearance and motion onset in this task (see **Supplement 1** for more information).

### 3.2. Dose-Testing: Behavioral Results

Using performance accuracy data from the 14 participants who completed both experimental sessions, the 2 (TMS Timings) X 5 (Stimulation Conditions) rmANOVA revealed a main effect of Stimulation Timing (*F*[1,13] = 5.754, *p* = .032, η^2^=0.307), with higher accuracy for N2–Onset, relative to Pre–Onset. The effect is illustrated in **Figure 3A**. In addition, the TMS Timings by Stimulation Conditions interaction was significant (*F*[4,52] = 3.313, *p* = .017, η^2^ = 0.203). Pairwise comparison results demonstrated that, when the stimulation was delivered at the timing of N2-onset, the condition of 80% at V5 led to higher response accuracy than 120% at V5 and 120% at Vertex (*ps* ≤ .042).

**Figure 3.**
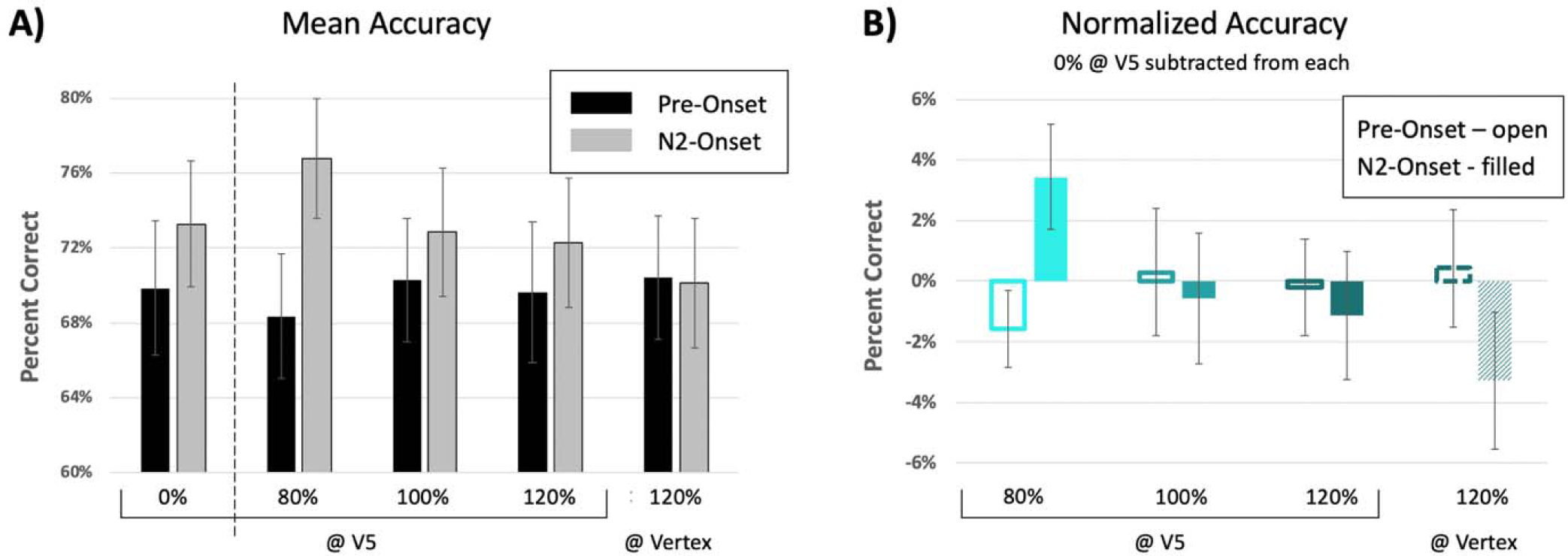
A) Mean accuracy for the five stimulation conditions, for the Pre-Onset (black) and N2-Onset (grey) TMS Timings. ANOVA showed significantly higher accuracy for N2-Onset and a significant interaction with higher accuracy at 80% intensity. B) Adjusted mean accuracy for each Stimulation Condition and TMS Timing minus the 0% condition, shows a significant interaction. Different shades indicate different intensity conditions with lighter colors indicating lower intensities.

Performance for each intensity level was also adjusted relative to the 0% at V5 separately for timing condition (e.g., Accuracy_80%_ - Accuracy_0%_) as a means to control for ongoing tonic variability in behavior in the absence of sound or magnetic stimulation elicited by the pulse. rmANOVA results on the adjusted behavioral accuracy, shown in **Figure 3B**, demonstrated non-significant main effects of TMS Timing (*p* = .944) and Stimulation Condition (*p* = .360) but a significant interaction (*F*[3,39] = 4.073, *p* =.013, η^2^ = 0.239). Pairwise comparison results showed that, when the stimulation was delivered at the timing of N2-Onset, the 80% at V5 condition led to higher response accuracy than 120% intensity at both V5 and Vertex locations (*ps* ≤ .046).

### 3.3. Dose-Testing: EEG Results

#### 3.3.1. Pre-Onset Stimulation Timing

Similar to the null results (*p* = .90) in behavioral evidence, one-way ANOVA results showed that N2 VEP amplitudes (i.e., early and late N2 responses in the V5 ROI) were not responsive to different Stimulus Conditions (*p* .314). In contrast, significant differences of P3 VEP amplitude were observed among different Stimulation Conditions (*F*[2.222,37.78] = 4.697, *p* = .013, *η*^2^ = 0.216). Pairwise comparison results showed that, as displayed in **Figure 4A**, mean P3 amplitude for the adjusted 80% at V5 condition was significantly smaller than those of all the other TMS conditions (*p*s ≤ .014). To further explore the relationship between behavioral accuracy and VEP changes, Pearson correlations were estimated between each of the three adjusted VEP amplitudes (i.e., early N2, late N2, and P3) and the adjusted response accuracy across all 18 participants. None of the correlation estimates reached statistical significance with corrected alpha level (i.e., .05/3 = .017).

**Figure 4.**
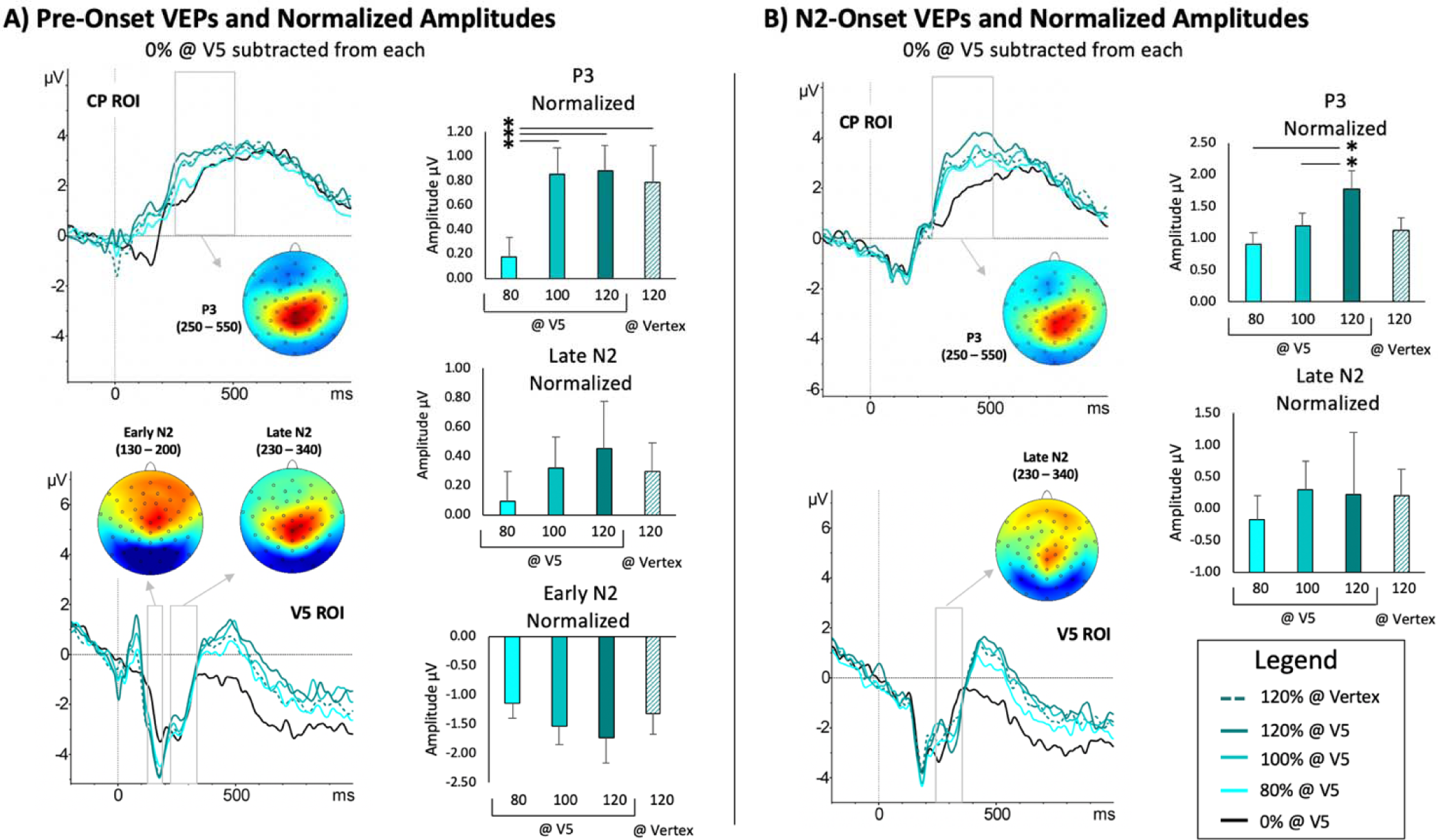
A) Pre-Onset VEPs, time-locked to the start of motion and shown for the CP (top) and V5 (bottom) ROIs for each of the five Stimulation Conditions (legend in bottom right). CSD topographies for the early N2, late N2 and P3 components are shown in relation to the grey boxes highlighting the latencies that are considered. Bar graphs represent the normalized (subtracting the condition of 0% at V5) mean amplitudes for each Stimulation Condition and Component. B) N2-Onset VEPs, topographies and normalized amplitudes in the same format as panel A, but without the early N2 component, which is obscured by TMS artifact.

#### 3.3.2. N2-Onset Timing

One-way ANOVA performed on the four normalized behavioral accuracy scores, revealed significant differences between conditions (*F*[3,42] = 4.794, *p* = .006, *η*^2^ = 0.255), with higher accuracy for the 80% RMT condition relative to all the other conditions (*ps* ≤ .032), confirming the behavioral effect in this sub-sample of data.

One-way ANOVA results on the late N2 amplitude in the V5 ROI and the P3 amplitude in the CP ROI^1^ revealed significant main effects of Stimulus Condition on the P3 amplitude (*F*[3,42] = 4.475, *p* = .008, *η*^2^ = 0.242), but not in the late N2 component (*p* = .711). Pairwise comparisons indicated that the 120% at V5 condition resulted in higher mean amplitude than the 80% (*p* = .003) and 100% (*p* = .019) conditions, as illustrated in **Figure 4B**. The Pearson correlation between adjusted VEP amplitudes and adjusted response accuracy across individuals were explored similarly to those in the Pre-Onset timing. No significant correlation estimates were identified for either the late N2 or P3 components with corrected alpha level.

## 4. Discussion

The current study aimed to identify task-relevant dose–response functions of single pulse TMS in order to examine the influence of stimulation timing and intensity on electrophysiological and behavioral responses. Here, single pulse TMS was applied at two latencies relative to the onset of psychometrically-calibrated, near-threshold, motion stimuli to assess the behavioral and electrophysiological changes due to TMS. It was hypothesized that stimulation would lead to greater behavioral disruption in motion direction discrimination when stimulated at the onset of the N2 component, given the critical links between this component and motion perception, and the past reports of acute perceptual disruption due to online TMS (Beynel et al., 2019). This was expected to interact with stimulus intensities such that higher intensities would induce greater behavioral effects. It was observed that stimulation applied at the N2-Onset led to behavioral facilitation in motion discrimination relative to Pre-Onset stimulation, however, this effect interacted with stimulation intensity revealing greater facilitation of motion discrimination accuracy at the lowest stimulation intensity of 80% RMT. Furthermore, it was found that VEP amplitudes significantly differed for the P3 component, with lower amplitudes in the 80% intensity condition for the Pre-Onset condition and higher amplitudes in the 120% intensity condition for the N2-Onset condition. These results suggest that timing and intensity interact with the profiles of perceptual and electrophysiological responses to near threshold motion stimuli with the additional indication that behavioral facilitation is being promoted by lower intensity stimulation.

### TMS effects on motion perception

Past studies have reported a wide variety of behavioral effects that result from single or paired pulse TMS over a range of factors, such as timing, intensity, and the ongoing activation state of the stimulated region (de Graaf et al., 2014; Romei, Thut, & Silvanto, 2016; Sandrini, Umiltà, & Rusconi, 2011; Wagner, Valero-Cabre, & Pascual-Leone, 2007). Although single-pulse TMS has mainly been shown to be disruptive (Abrahamyan, Clifford, Arabzadeh, & Harris, 2015; Amassian et al., 1989; Beckers & Zeki, 1995; Desmond, Chen, & Shieh, 2005), it has been reported that facilitation can be achieved with the right combination of stimulation timing, intensity and background brain state (Abrahamyan, Clifford, Arabzadeh, & Harris, 2011; Abrahamyan et al., 2015; Silvanto et al., 2017; Silvanto, Bona, Marelli, & Cattaneo, 2018). For example, enhanced target detection has been found after stimulating the visual cortex with low intensity single pulse TMS delivered 100 and 120 ms after stimulus onset (Abrahamyan et al., 2011, 2015). In these cases, it was proposed that the summation of ongoing neural responses and the excitation induced by the lower intensity TMS pulse resulted in enhanced processing and behavioral facilitation. This explanation is similar to the stochastic resonance account (Schwarzkopf, Silvanto, & Rees, 2011; Stocks, 2000) which proposes that the strength of a stimulus can be improved by externally enhancing the ongoing neural activity. That is, a single pulse of TMS, delivered at low intensity, will add low levels of noise to the neural system, boosting information exchange between the neurons in the stimulated cortex (Silvanto et al., 2017).

While such a model may explain the observed effects, other factors may also contribute to facilitation or inhibition in such contexts (Cattaneo & Silvanto, 2008). For example, adaptation that may result from repeated presentations of a stimulus has been shown to influence TMS effects with demonstrations that TMS can reverse the influence of adaptation on target detection (Silvanto, Muggleton, Cowey, & Walsh, 2007). Similarly, TMS applied during the delay between a visual priming stimulus and a target stimulus also leads to selective modulation of the target to reduce the priming effect, demonstrating that attributes encoded by less active neural populations are preferentially facilitated by TMS (Cattaneo & Silvanto, 2008). Finally, as proposed in other studies demonstrating modulatory effects of TMS to V5 in the N2 latency range (Laycock et al., 2007; Sack et al., 2006), may reflect feedforward/feedback processing between the striate and extrastriate cortices. The current findings build on these theorized mechanisms to provide additional insight into state-dependency TMS effects by describing systematic dose influences on behavior, and particularly the observed profile with greater enhancement effects at lower intensities.

### TMS effects on motion VEPs

The combination of transcranial magnetic stimulation and electroencephalography (EEG) is a powerful tool for investigating cortical mechanisms and networks with fine temporal precision. Using this technique, we expected to obtain electrophysiological evidence of the interplay between brain activity and TMS conditions. Our results showed that the mean activity of the P3 component in the CP ROI produced significant differences between intensities in both timing conditions. In general, high intensity single pulse TMS evoked higher mean activity values for the P3 component than pulses at low intensities. In the Pre-Onset 80% condition, values were significantly smaller than for the higher intensity TMS conditions, while values were significantly higher for the 120% condition at N2-Onset when compared to the lower intensity conditions, but not to vertex-120%. This mean activity increase, however, did not correlate with the magnitude of behavioral effects across participants. While N2 amplitudes generally scaled monotonically with higher stimulation intensities, these did not differ significantly across conditions or correlate with behavioral effects.

In interpreting these effects, it is possible that the EEG response obtained here may have been impacted by indirect brain stimulation. Namely, despite the presence of both a no-stimulation control and a vertex control, the lack of a somatosensory-matched sham condition is a limitation of this study and the observed intensity effects may be attributed to (multi)sensory evoked activity and not necessarily the result of neural changes due to the induced magnetic field (Casarotto et al., 2010; Conde et al., 2019). As discussed by others (Siebner, Conde, Tomasevic, Thielscher, & Bergmann, 2019), an important contributor to the TMS-EEG response is the prominent auditory evoked potential associated with the clicking sound produced by TMS. This auditory response is introduced by a combination of bone-conduction and air conducted sound reaching the cochlea and activating central and parieto-temporal regions bilaterally to produce a prominent N1-P2 complex (Lioumis, Kičić, Savolainen, Mäkelä, & Kähkönen, 2009). Reducing sound levels and distancing the coil from the head, have been shown to reduce the N1-P2 complex amplitude (Seppo Kähkönen, Wilenius, Komssi, & Ilmoniemi, 2004; Nikouline, Ruohonen, & Ilmoniemi, 1999; ter Braack, de Vos, & van Putten, 2015), and while earplugs and foam insulation were used to mitigate sound here, it is possible that these measures were not enough to suppress the auditory response. Thus, future studies should strive to use a more sensitive somatosensory-matched sham control condition to better isolate TMS effects. An alternative explanation for the amplitude scaling of the P3 response, but not the N2 response may pertain to attentional capture that is greater for higher intensity stimulation. As noted, the P3 response is strongly associated with attentional processes (Kuba et al., 1998; Kubová et al., 2002) and therefore, a plausible explanation of the effects here may stem from uncontrolled attentional allocation.

An additional, related observation in this study was that during the Pre-Onset condition we found the presence of an early component at a latency roughly equivalent to the P1. This response was not present in the 0% intensity condition, nor during the early stages of the EEG response in the N2–Onset condition, pointing to the possibility that this response may have been driven entirely by the TMS pulse and not associated with the visual stimulus or the interaction between visual and magnetic stimulation. Further control sessions, performed with a subset of seven participants, who received TMS without visual stimulation (see **Supplementary Figure 1**) also produce a robust EEG response in the P1 range, further supporting this assumption that this response is driven by the TMS pulse rather than interactions with evoked visual responses. As noted above, such findings need to be considered in the context of multisensory stimulation and associated artifacts in the TMS response (Bonato, Miniussi, & Rossini, 2006; S. Kähkönen, Komssi, Wilenius, & Ilmoniemi, 2005; Lioumis et al., 2009).

### Future directions and conclusions

While the present study offered a unique view of the dose-response relationships between TMS intensity, behavior and electrophysiological brain responses, there are a number of limitations that should be improved upon in future studies. As noted in the section above, one such limitation is possible presence of multisensory stimulation confounds that should be better controlled using somatosensory-matched electrical stimulation in future studies. Another important consideration is the mode used to define stimulation intensity, which here, was delivered relative to resting motor threshold. While such intensity scaling is principled, objective, and widely used, this dosing is made based upon motor system responsiveness and not visual system responsiveness. The current finding therefore need to be considered in light of previous research showing that MEPs can be elicited at intensities below RMT, and that in motor cortex there are distinct recruitment (i.e. input-output) curves for inhibitory processes that typically elicit lower threshold than for excitatory processes (Kallioniemi, Säisänen, Könönen, Awiszus, & Julkunen, 2014). Moreover, while there is also a literature showing that phosphene thresholds are generally higher than motor thresholds (Antal, Nitsche, Kincses, Lampe, & Paulus, 2004; Boroojerdi et al., 2002; Deblieck, Thompson, Iacoboni, & Wu, 2008), there is also considerable heterogeneity in ability to elicit phosphenes in individual subjects. In light of these challenges, future studies should build on the current dose-response functions to include intensity scaling based on visual responsiveness. Lastly, it is worth noting that there is an important movement towards larger and better powered sample sizes in TMS studies. While the current sample of 15 individuals who completed all study activities is larger than average sample size of 11, reported in a recent meta-analysis of online TMS studies published last year (Beynel et al., 2019), future studies should continue to strive for even larger sample sizes.

## Acknowledgments

The authors would like to thank Zachary Abzug, Lysianne Beynel, Tracy D’Arbeloff, Erik A. Wing, and Rachel Donaldson, who helped develop the behavioral task and Lari Koponen who help with the concurrent TMS-EEG protocols used in this research. The authors would also like to thank all of the individuals who participated in this research.

## Supplement

**Supplementary Figure 1.**
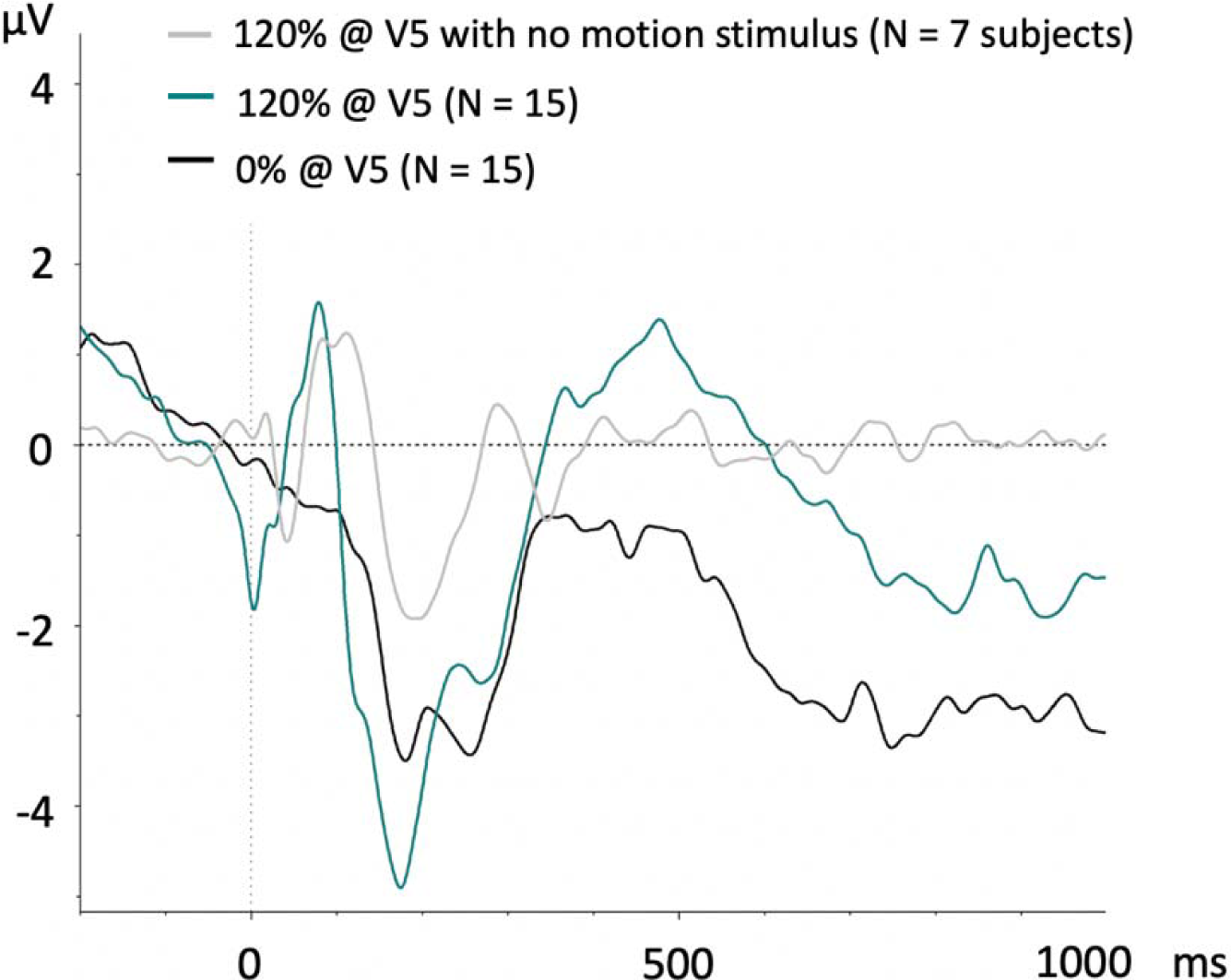
In addition to the visual evoked potentials collected as part of the main experiment, seven participants were also tested on a brief supplemental session in which 120% RMT stimulation was delivered over their V5 target as they fixated on a screen with no visual motion stimulus. Here, 120 trials of single pulse TMS was delivered at random intervals spaced apart between 7 and 10 seconds as they gazed at an unchanging fixation mark To illustrate the difference in evoked responses to TMS delivered in the presence of visual motion (replicated from **Figure 4A** bottom), and responses in the absence of visual motion, these VEPs are superimposed in **Supplemental Figure 1**. Here evoked responses from 120% stimulation with motion stimuli (green trace) and 120% stimulation with no motion stimuli (grey trace) both elicited a pronounced positive component in the range of 100-120 ms. Conversely, VEPs elicited to the onset of motion, but in the absence of single pulse TMS (black trace) did not produce such a positive P1 component.

## Footnotes

1 the early N2 component could not be analyzed because it overlapped with the TMS pulses and was removed during data cleaning

